# Brain Stimulation Prevents Neural Downregulation and optimizes Learning

**DOI:** 10.1101/2024.11.06.622343

**Authors:** F. Contò, G. Ellena, G. Edwards, L. Battelli

**Affiliations:** Center for Neuroscience and Cognitive Systems@UniTn, Istituto Italiano di Tecnologia, Corso Bettini 31, 38068 Rovereto (TN), Italy; National Institutes of Health, Bethesda, Maryland 20892, USA; Berenson-Allen Center for Noninvasive Brain Stimulation, Department of Neurology, Beth Israel Deaconess Medical Center, Harvard Medical School, Boston, MA 02215

**Author notes:** **Corresponding Author:** Federica Contò. **Competing financial interests:** The authors declare no competing financial interests.

## Abstract

Non-invasive brain stimulation, such as transcranial random noise stimulation (tRNS), has been shown to enhance cortical excitability and facilitate perceptual learning. However, the neural mechanisms underlying these effects remain poorly understood. Here, we demonstrate that tRNS over the bilateral intraparietal sulcus (IPS) optimizes learning by preventing the decline in neural activity that occurs during short, high-load attentional training, thereby sustaining excitability and enhancing behavioral performance. Using a multi-session tRNS-fMRI paradigm, we investigated how tRNS modulates learning-related plasticity in the attention network during cognitive training. In the sham condition we observed a significant decline in task-evoked BOLD after the training and no behavioral improvement, suggesting crucial neural changes associated with cognitive training that are not evident in the behavioral data. Conversely, in the active tRNS condition, stimulation not only prevented the early decline in task-evoked BOLD activity observed in the sham condition, but it increased it. This change in BOLD activity correlated with improved performance. This suggests that tRNS counteracts early neural adaptation during short training protocols, sustaining activity in task-relevant cortical regions to enable learning that would otherwise fail. By providing the first direct evidence that tRNS mitigates early neural downregulation and preserves functional response dynamics during learning in crucial task-related cortical areas, our study demonstrates that in parietal cortex training-induced plasticity is not accompanied by efficiency-driven reductions in activation, like typically seen in sensory areas. Instead, we propose that sustaining neural excitability through tRNS prolongs plasticity and optimizes cognitive performance in higher order attentional areas. These findings highlight tRNS as a powerful tool for enhancing attentional learning and modulating neuroplasticity in both healthy and clinical populations.

## Introduction

Perceptual learning (PL) refers to the sustained improvement of sensory abilities through experience and practice ([1], [2][3]). This process, along with the associated neural plasticity [4], [5], [6], [7], underscores the brain’s remarkable adaptability and plays a crucial role in navigating complex environments. PL also has significant implications for developing interventions that enhance sensory perception and cognition. Attentional processes are deeply intertwined with PL, with the dorsal-ventral cortical attention network ([8], [9], [10]) playing a key role in guiding perceptual training. Attention selectively enhances relevant sensory inputs ([11], [12], [13], [14], [15], [16], [17], [18]), and fMRI studies have shown that distinct activation patterns in the dorsal and ventral attentional networks correspond to different aspects of learning ([7], [11][19]). Despite evidence linking attention, learning, and neural plasticity, the precise interplay between these mechanisms remains unclear.

While extensive training often results in robust performance improvements ([3]), shorter training sessions frequently fail to induce lasting learning, particularly under high attentional demands. This failure has been attributed to a rapid decline in neural excitability within task-relevant cortical regions ([20]), such as the intraparietal sulcus (IPS), a key hub of the dorsal/ventral attention network (DVAN) involved in top-down control ([21], [22], [23], [24]). fMRI studies suggest that successful learning is associated with sustained IPS activity, whereas unsuccessful learning is marked by an early decline in activation, likely due to efficiency-driven stabilization of neural representations ([22], [23], [25], [26]). Neurophysiological studies further indicate that PL involves both enhanced sensory tuning and strengthened cortico-cortical communication ([27]). Long-term training sharpens neuronal selectivity within sensory regions ([28]) while reinforcing connectivity with higher-order areas like the IPS, which integrate sensory evidence and guide decision-making ([3], [27], [29]). PL is not static; rather, the brain regions mediating learning shift dynamically with training ([30]). This shift is supported by evidence that PL engages widespread networks beyond early visual areas, including higher-order parietal and decision-making regions ([31], [32]). For example, Chen et al. ([27]) found that motion discrimination training not only sharpened stimulus-specific tuning in V3A but also optimized connectivity with IPS, reinforcing the idea that parietal regions play a critical role in integrating sensory and decision-related information. In the non-human primate literature, the lateral intraparietal area (LIP), homologous to the human IPS, exhibits learning-related response changes ([29]). Notably, for instance, face perception training selectively enhances plasticity in the left fusiform cortex ([33]), emphasizing that PL mechanisms are task-dependent and engage distinct brain circuits based on the nature of the learned feature. However, during brief training, these processes may not fully develop, leading to early downregulation of IPS activity and learning failure ([23]). Understanding how to sustain neural plasticity during early learning phases is therefore critical to optimizing training outcomes ([29]).

Transcranial random noise stimulation (tRNS), a non-invasive brain stimulation technique, has been proposed to enhance PL by modulating cortical excitability and preventing premature stabilization of neural activity ([7], [34], [35], [36]). Studies suggest that tRNS facilitates PL across multiple domains, particularly when applied over visual and parietal regions ([7], [14], [37], [38]). However, its mechanisms in attention-dependent learning remain poorly understood. This study examines how tRNS over the bilateral IPS influences plasticity during short, high-load attentional training. Using a multi-session tRNS-fMRI paradigm, we test whether tRNS prevents the early decline in task-evoked IPS activity associated with unsuccessful learning ([23];[20]). Specifically, we analyze BOLD responses during two attentional tasks (an orientation discrimination and a temporal order judgment task) to assess how stimulation influences neural plasticity. We hypothesized that IPS stimulation will prolong early-stage plasticity and enhance learning rates compared to sham stimulation.

Our findings reveal that tRNS sustains neural excitability in the IPS, preventing the BOLD activity decline seen in the sham condition while improving behavioral performance, as shown in our previous work ([7]). These results challenge the notion that efficiency-driven reductions in cortical activity are necessary for learning and instead underscore the importance of maintaining activation in attentional control regions ([25], [30]). By demonstrating that tRNS counteracts early neural adaptation and preserves functional response dynamics, our study provides novel insights into how non-invasive stimulation facilitates PL and cortical plasticity. These findings have broad implications for both fundamental neuroscience and clinical interventions aimed at enhancing cognitive function in healthy and impaired populations ([3]).

## Results

We conducted task-related fMRI analysis to investigate the impact of multi-session tRNS combined with training on long-term modulation of the BOLD signal during two tasks: an orientation discrimination (OD) and a temporal order judgment (TOJ) task. Participants underwent fMRI scanning during both pre- and post-test sessions (S2 and S7, *Figure 4*; *Methods* section), during which they performed the two attentional tasks (see *Methods* section for details on the tasks). In the training phase, which took place between the two fMRI sessions, all subjects were asked to perform two tasks simultaneously, alternating on each trial, and were assigned to three different groups. One group received tRNS over the bilateral IPS, our crucial hotspot for visuospatial attention ([8], [39]) and a central hub for the DVAN ([40]). A second group received tRNS over the bilateral human middle temporal (hMT) areas (active control condition) and a third group received Sham stimulation. Of note, the active control condition was particularly challenging given the close proximity of the hMT to the IPS, but it was important to test both areas as they are part of the DVAN but may contribute differently to visuospatial processing and thus may respond differently to stimulation and training.

To evaluate the impact of multi-session tRNS combined with task training, we measured changes in the BOLD signal between pre- and post-test session for each task individually. Single-subject deconvolution analysis was performed to examine the BOLD responses between the two different stimuli conditions (orientation and temporal stimuli). Peaks of the individual deconvolved beta weights relative to the ROIs embedded in the DVAN were analyzed to evaluate the impact of stimulation combined with training on the attentional network. In addition, to control for network-specific effects of stimulation, we also investigated whether tRNS and training impacted cortical activity within a different (control) network, the default mode network (DMN), which is typically active during passive or resting conditions ([41], [42], [43], [44]). Next, we compared the change in task-evoked beta weights measured within the DVAN and the DMN to test whether these two networks showed different modulation of the BOLD activity measured within their main ROIs after stimulation. Finally, we investigated the change in beta weights of each single ROI embedded in the attentional network across different conditions. Details of data analysis pipeline can be found in the Methods section.

### Impact of stimulation on the DVAN

We first ensured that the three stimulation groups did not differ in brain activity at baseline by comparing their beta weights on the pre-test fMRI session for each task. To do so, we fitted a linear mixed effects model (LMM) to examine the effect of different stimulation conditions on the normalized beta weights measured during the pre-test, while accounting for individual and ROIs differences (the model included random intercepts for subjects and for brain areas). The Type II Wald Chi-Square test for the Stimulation Condition effect factor yielded a Chi-square value of 4.102 (χ^2^(2) =4.102, *p* = 0.129, *lmer*) for the OD task and 1.717 (χ^2^(2) =1.717, *p* = .423, *lmer)* for the TOJ task, indicating that the overall effect of different stimulation conditions on beta weights was not statistically significant at baseline for either task.

Next, delta scores representing the difference between pre- and post-stimulation beta weights were computed at the individual subject level (Δβ = β(fMRI post-test) – β(fMRI pre-test) and per single ROI to quantify changes in BOLD signal modulation. These ROIs include key areas such as the anterior, posterior, and central intraparietal sulcus (IPS), human middle temporal (hMT), frontal eye fields (FEFs), temporal parietal junction (TPJ), ventral frontal cortex (VFC), and inferior frontal junction (IFJ), covering both dorsal and ventral attention networks (see *Methods* section for a complete list of the ROIs included in the DVAN, and the Talairach coordinates). The delta values indicate the changes in BOLD signal modulation normalized to baseline activity, before stimulation. An LMM was then fitted to examine the combination of stimulation and training on subjects’ delta beta weights (normalized delta beta weights were modeled as a function of Stimulation conditions, with random intercepts for subjects and brain areas). This analysis revealed a main effect of stimulation group on delta beta weights in response to the OD task (χ^2^(2) =12.787, *p* =.0017, *lmer)*, indicating that changes in beta weights were significantly modulated by the type of stimulation received during training (*Figure 1*). We then compared delta values between different stimulation conditions, and found a significant difference between the Parietal and Sham group with an estimate of 0.678 (SE=0.193, p=.0041 FDR-adjusted), and between the Parietal and hMT group with an estimate of -0.442 (SE=0.189, p=0.039 FDR-adjusted), indicating that the Parietal condition resulted in significantly higher delta beta weights compared to the Sham and hMT condition (*Figure 1*). No significant difference was found when comparing delta values of the hMT and Sham condition (estimate of 0.236; SE=.189; *p*=.2207 FDR adjusted), indicating that the difference of delta beta weights was not modulated differently between these two conditions. Further analysis of the estimated marginal means (EMMeans), used to summarize the effects of different factors in the linear model, revealed that the average delta values for the Parietal condition was of 0.335 (SE=0.141, with a 95% CI of 0.049 and 0.621), while for the Sham the average delta beta weights was of -0.343 (SE=0.141, with a 95% CI of - 0.63 and -0.057), and for the hMT condition of -0.107 (SE=.135, with a 95% CI of -.383 and .0167). These data suggest that the delta beta weights increased significantly after parietal stimulation, while they decreased significantly in the Sham condition. Finally, the delta beta weights of the DVAN during the OD task did not change in the hMT stimulation condition.

**Figure 1.**
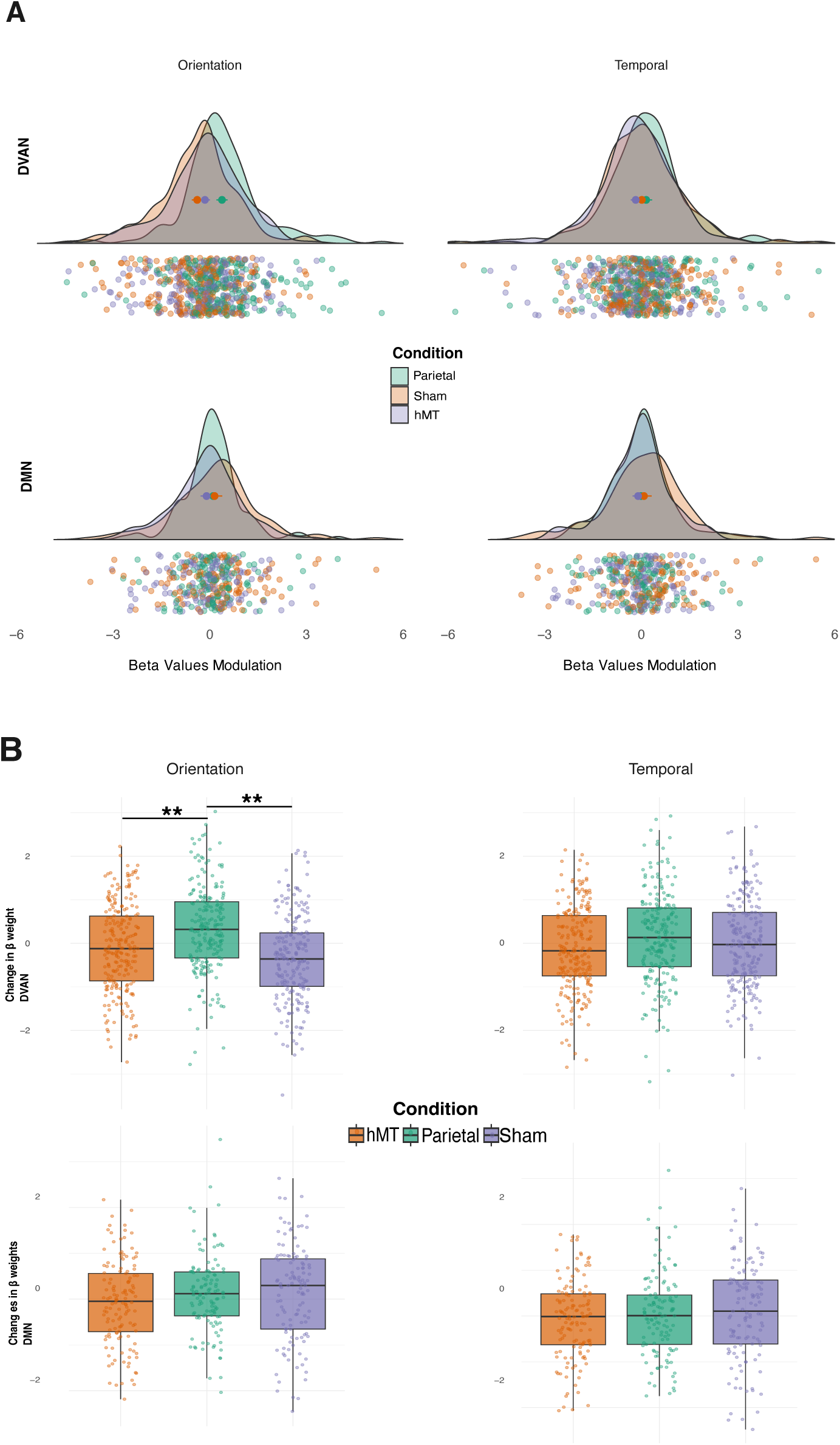
Modulation of Beta weights across different stimulation conditions and networks in response to the two attention tasks (OD and TOJ). A. Change of Beta weights frequency distribution for each stimulation condition computed as the difference between beta weights pre- and post-test session (Δβ = β(fMRI post-test)− β(fMRI pre-test)); orange for hMT, green for Parietal and violet for Sham). Single bold Dots within distributions indicated the median for each stimulation condition, while dots below the distribution curves represents individual data measured for each participant in each of the ROIs embedded in the two networks (Top panel: DVAN, bottom panel: DMN) in response to each type of task stimuli (Orientation on the left and Temporal on the right). B. Box plots for β-weights changes by stimulation condition in response to the OD (left Panel) and the TOJ task (right Panel) normalized to baseline, prior to stimulation. The center line in the middle of the box is the median of each data distribution, while the box represents the interquartile range (IQR), with the lower quartile representing the 25th percentile (Q1) and the upper quartile representing the 75th percentile (Q3). Dots represents individual subjects’ data. Lines beyond the box represent the minimum and maximum values in the data. Asterisks indicate significant differences between two stimulation condition.

We then analyzed the effect of different stimulation conditions on the normalized delta beta weights measured in the ROIs of the DVAN during the temporal task (TOJ) using a similar LMM as for the OD task (normalized delta beta weights relative to the TOJ task were modeled as a function of Stimulation conditions, with random intercepts for subjects and brain areas). The Type II Wald Chi-Square test revealed no significant effect of stimulation condition on the normalized delta beta weights (χ^2^(2) =1.917, *p* =.3834, *lmer*), indicating that the type of stimulation condition did not significantly influence the changes in beta weights for the TOJ task. Pairwise comparison run following the model between conditions showed no significant differences between any pair of conditions (hMT vs Parietal comparison estimate of -0.274, SE=.198, *p*=.361 FDR-adjusted; Parietal vs Sham comparison estimate of 0.14, SE=.202, *p*=.768 FDR-adjusted; hMT vs Sham comparison estimate of -0.134, SE=0.198, *p*=.779 FDR-adjusted).

Next, to control for network-specific effects of stimulation, we analyzed changes in BOLD activity within the DMN, which we hypothesized would not to be affected by the stimulation protocol or by the training phase. The same pipeline analysis for the DVAN was applied to the DMN. Therefore, we first fitted an LMM to examine whether there was a difference in beta weights at baseline across the three stimulation groups, with random factors being subjects and brain areas. No significant main effect was found for the OD task (χ^2^(2) =0.967, *p* = .616, *lmer*), nor for the TOJ task (χ^2^(2) =0.357, *p* = .836, *lmer)*, indicating that beta weights measured at baseline did not differ between the three stimulation groups in this control network. Next, as for the DVAN, delta scores were calculated for each ROI of the DMN to quantify changes in BOLD signal modulation (Δβ = β(fMRI post-test) – β(fMRI pre-test) and were used to fit an LMM that examined the effect of stimulation and training on delta beta weights changes, with random intercepts for subjects and ROIs. No significant main effect of stimulation condition was found on delta beta weights in response to the OD (χ^2^(2) =1.446, *p=.4851*, *lmer)*, or the TOJ task (χ^2^(2) =0.565, *p* =.7538, *lmer),* indicating that beta weights changes were not modulated differently by the type of stimulation received during training (*Figure 1*). Hence, none of the analyses conducted for the DMN revealed any significant effects of stimulation and training.

Finally, to investigate the specificity of the tRNS effect during training on attention tasks, we further explored differences in beta weight modulation across the three stimulation groups measured within the DVAN relative to DMN. An LMM was used to investigate the effect of stimulation combined with training (Parietal, hMT and Sham condition) and of network (DVAN and DMN) on normalized delta beta weights in response to the OD task, with a random effect for subjects. The analysis revealed significant effect of Condition (χ²=6.170, df=2, *p* = 0.046) and a highly significant interaction between Condition and Network (χ²=23.625, df =2, *p* < 0.001), while the main effect of network alone was not significant (χ²=2.633, df =1, *p* = 0.105). We then run post-hoc analysis using estimated marginal means (EMMs). Specifically, pairwise comparisons between networks within each condition indicated that the difference between modulation of beta weights was significant for the Parietal (estimate = -0.216, SE = 0.103, *p* = 0.036 FDR-corrected) and for the Sham condition (estimate = 0.482, SE = 0.103, *p* < 0.001 FDR-corrected), while no significant difference in beta weights modulation between the two networks was found for the hMT group (estimate = 0.026, SE = 0.099, *p* = 0.790 FDR-corrected). Differences in network-specific changes within each stimulation condition are depicted in *Figure 2* (top row).

**Figure 2.**
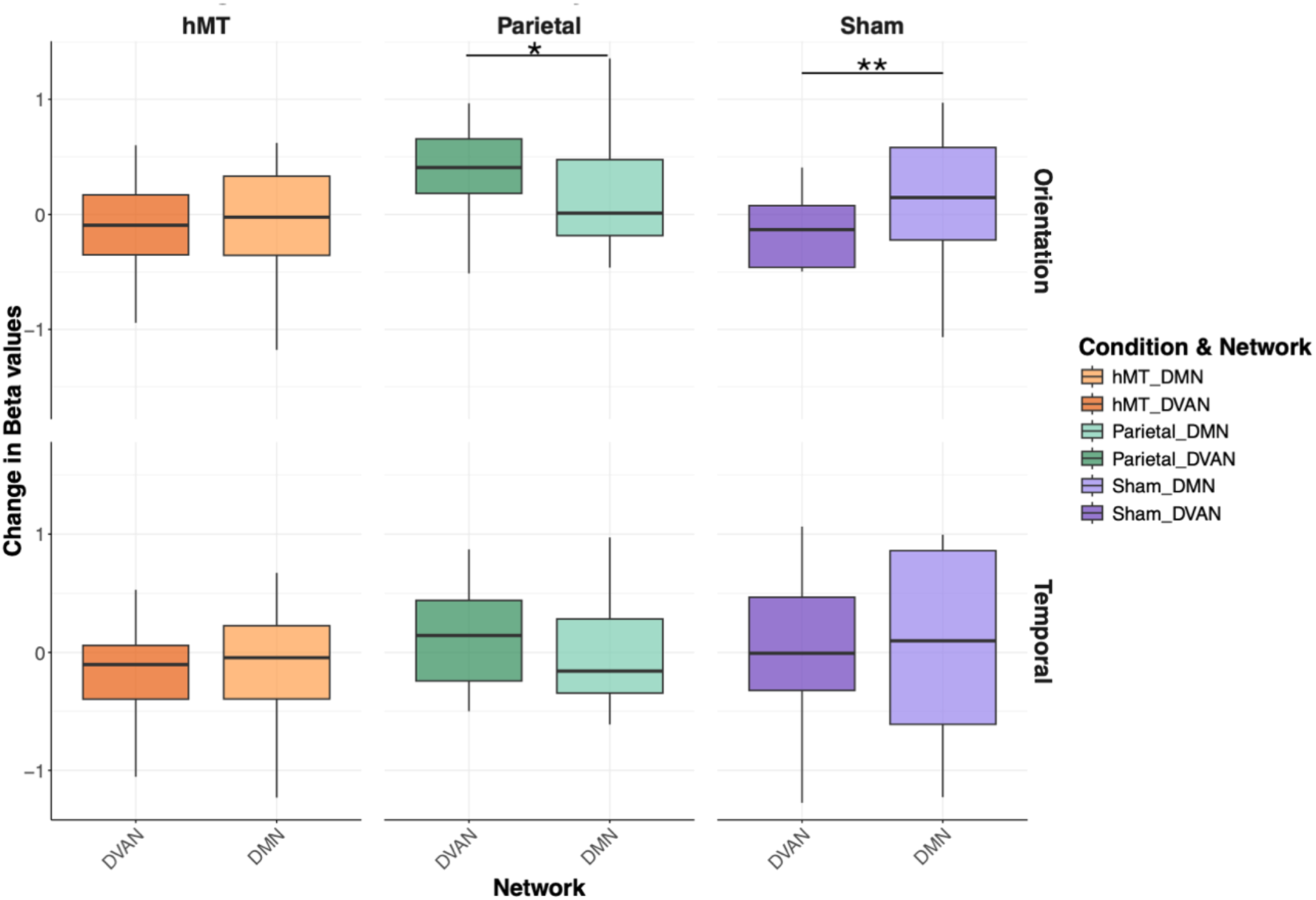
Box plots for Beta weights changes across conditions (hMT, Parietal and Sham) and networks (DVAN and DMN) in response to the OD (top row) and the TOJ task (bottom row) normalized to baseline and averaged across subjects per stimulation condition. The center line in the middle of the box is the median of each data distribution, while the box represents the interquartile range (IQR), with the lower quartile representing the 25th percentile (Q1) and the upper quartile representing the 75th percentile (Q3). Lines beyond the box represent the minimum and maximum values in the data.

Next, to further control for changes in the DMN relative to the DVAN, we ran the same LMM to analyze the beta weights relative to the TOJ task. In contrast to the finding for the OD, the results showed that there were no significant effects for Condition (χ²=1.341, df=2, *p* = 0.511), nor for network (χ²=.0004, df=1, *p* = 0.984), or for the interaction between Condition and Network (χ²=2.761, *p* = 0.251). These findings indicate that stimulation had no significant impact on the modulation of beta weights in response to the TOJ task for either the target network (DVAN) or the control network (DMN), nor the interaction between Condition and Network significantly influenced the normalized data (*Figure 2*, bottom row).

### Impact of stimulation on ROIs’ response across conditions after learning

Because the stimulation condition differently influenced the change in beta weights measured between pre and post-test session (expressed as delta) in response to the OD task at a network-level, we aimed to investigate whether stimulation condition exerted differential effects on different ROIs embedded in the DVAN. We first analyzed these effects across all regions of interest without the potential confounding factor of hemispheric lateralization that may affect the functional role of each ROI, potentially leading to differences in responsiveness to stimulation that are unrelated to the intended effects of the task (which is bilateral). An LMM was then fitted to examine the effects of stimulation on normalized delta values across different brain areas regardless of their lateralization, therefore data measured in the left and right hemisphere of each bilateral ROI were collapsed into one ROI. The results revealed significant differences in delta scores across stimulation conditions in several brain regions (*Figure 3A,* left Panel). Specifically, significant main effects of stimulation condition were observed in the hMT (χ²=19.892, df=2, *p* = <.001), anterior IPS (χ²=9.567, df=2, *p* = 0.008), posterior IPS (χ²=15.181, df=2, *p* =.0005), and FEF (χ²=6.280, df=2, *p* = .043) regions, while other regions did not show significant changes under different stimulation conditions (see *Table 1* below for detailed statistical results of all models run on each DVAN ROIs). Subsequent post-hoc analyses using estimated marginal means (EMM) further detailed these findings. Pairwise comparisons revealed significant differences between specific conditions in the hMT (Parietal vs. Sham: *p* <0.001 FDR-corrected, estimate=1.041, t.ratio=4.411; hMT vs. Sham: *p*=0.017 FDR-corrected, estimate=0.65, t.ratio=2.817), posterior IPS (Parietal vs. Sham: *p*<0.001 FDR-corrected, estimate=1.367, t.ratio=3.888) and anterior IPS regions (Parietal vs. Sham *p*=0.008 FDR-corrected, estimate =1.122, t.ratio=3.077), indicating robust effects of stimulation condition on delta beta weights for the OD task across these brain areas. All other pairwise comparisons were not significant. For the analysis of the anterior TPJ brain area, we could not fit an *lmer* model as for the other brain areas, because we had only one beta weight per subject (as this region is lateralized to the right); we then used an ANOVA, that was not significant (F(2,31)=0.224, *p*=.801).

**Figure 3.**
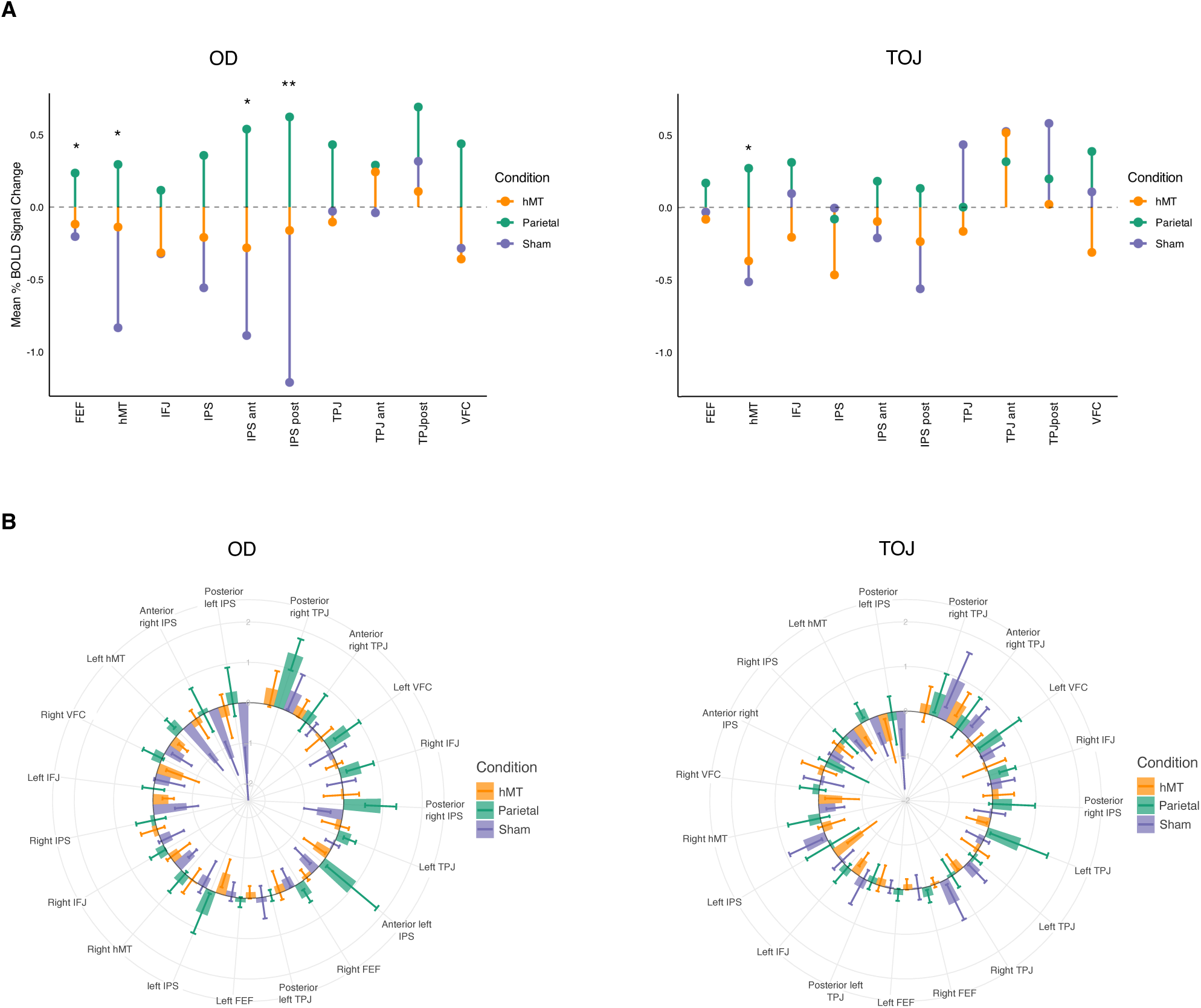
Beta weights changes per single ROI embedded in the DVAN. **A.** The lollipop chart illustrates the mean percentage BOLD signal change for each experimental condition and depicted across different brain areas (collapsed across hemisphere, x-axis). Each lollipop segment represents the mean value, lollipop segments above the dashed horizontal line at zero indicate increased BOLD signal, while segments below this line indicate decreased BOLD signal. The chart is facetted by Predictor (OD on the left and TOJ on the right), allowing for comparisons of stimulation conditions within each predictor, as well as between predictors. Asterisks indicate the ROI where there is a significant different BOLD modulation between two or more conditions. **B.** Changes between pre- and post-test BOLD signal is depicted with bar plots across Conditions (green for Parietal, orange for hMT and violet for Sham) and across tasks (OD on the left and TOJ on the right) per each ROI (data were differentiated per hemisphere). The line within each bar indicates the SEM.

**Table 1.** Results of the LMM for each ROI embedded in the DVAN, with data collapsed across hemispheres for the OD task. For each brain area (left and right hemisphere data were collapsed), statistical details of the LMM fitted (lmer(normalized Δβ ∼ Condition + (1|subj), data=df_ROI) are reported comprising of Chi-square and corresponding p-value of the model.

| ROI | ChiSq | Df | Pr(>Chisq) |
| --- | --- | --- | --- |
| FEF | 6.2806 | 2 | 0.0433 |
| hMT | 19.8925 | 2 | <.0001 |
| IFJ | 3.0996 | 2 | 0.2123 |
| IPS | 4.5595 | 2 | 0.1023 |
| Ant IPS | 9.5672 | 2 | 0.0084 |
| Post IPS | 15.1813 | 2 | 0.0005 |
| TPJ | 3.9429 | 2 | 0.1393 |
| Post TPJ | 1.7965 | 2 | 0.4073 |
| VFC | 4.5445 | 2 | 0.1031 |

Next, we further investigated whether the effects observed are consistent across the network with an analysis for each ROI divided per hemisphere. As every single brain area (divided per side) was analyzed independently, and no mixed-linear effects model could be fitted, EMMeans was used to run a series of pairwise comparisons (on delta beta weights) between the stimulation conditions, this time for each of the ROI included in the attentional network divided per hemisphere (*Figure 3B*). This analysis provides estimated marginal means for each combination of the factors “Stimulation” and “ROI”, along with confidence intervals and standard errors, and it allows us to compare the estimated means between different conditions within each level of brain area (ROI), divided per side. A difference in the change of the beta weights relative to the OD task was found in the following brain areas: left hMT (parietal vs sham, estimate = 1.378, t.ratio=3.242, *p*=.004 FDR-corrected); left anterior IPS (parietal vs sham, estimate = 1.062, t.ratio=2.498, *p*=0.024 FDR-corrected; Parietal vs hMT estimate -1.0075, t.ratio=-2.422, *p*=0.024 FDR-corrected); right anterior IPS (Parietal vs Sham, estimate=1.183, t.ratio=2.782, *p*=0.017 FDR-corrected), left posterior IPS (Parietal vs Sham, estimate=1.465, t.ratio=3.447, *p*=0.0019 FDR-corrected), and right posterior IPS (Parietal vs Sham, estimate=1.268, t.ratio=2.984, *p*=0.009, FDR-corrected). No significant difference between any pair of conditions was found for the other ROIs of the attention network. A complete report of all the analysis comparing the delta values for each ROIs divided per hemisphere is reported in the Appendix (*Table 1*, Appendix). These results indicate that the stimulation condition has a significant and region-specific impact on neural responses during the OD task, revealing distinct activation patterns under different experimental conditions.

To understand the individual ROI responses to both tasks, we conducted the same analysis on data collected in response to the TOJ task. Consistent with the absence of significance in the main network model, no differences were found between stimulation conditions for any of the ROIs included in the attention network, whether the data was collapsed across hemispheres (*Figure 3A*, right panel) or not (*Figure 3B*, right panel), except for the hMT area, which showed a significant effect only when data were collapsed across hemispheres (χ²=8.29, df=2, *p* =0.016). The subsequent pairwise comparison showed a significant increase in BOLD signal within the hMT ROI in comparison to the Sham condition, in the Parietal group only (estimate =0.7, t-ratio= 2.69, p=0.024 FDR-corrected). A complete report of the results relative to the TOJ task are reported in the Appendix (*Table 2* and *Table 3*). Taken together, these results reveal different patterns of activation depending on the specific experimental conditions used, indicating that the stimulation condition significantly affects the neural responses to the two attention tasks differently in different brain regions.

**Table 2.** Regions of Interest Coordinates. Standardized Talairach coordinates (X,Y,Z) for the centroid of each region of interest (Left and Right hemisphere) for the DVAN.

|  | LEFT HEMISPHERE |  |  | RIGHT HEMISPHERE |  |  |
| --- | --- | --- | --- | --- | --- | --- |
|  | X | Y | Z | X | Y | Z |
| IPS | -26 | -63 | 48 | 25 | -60 | 48 |
| Anterior IPS | -25 | -55 | 50 | 25 | -59 | 56 |
| Posterior IPS | -19 | -63 | 60 | 17 | -65 | 54 |
| hMT | -44 | -67 | 3 | 44 | -71 | 6 |
| FEF | -30 | -5 | 48 | 30 | -5 | 48 |
| TPJ | -48 | -54 | 26 | 48 | -54 | 21 |
| VFC | -42 | 12 | -1 | 40 | 17 | 4 |
| Anterior TPJ |  |  |  | 54 | -41 | 28 |
| Posterior TPJ | -44 | -66 | 22 | 49 | -61 | 20 |
| IFJ | -40 | 7 | 25 | 44 | 12 | 27 |

### BOLD changes and Behavior Correlation Analysis

Given the significant effect observed in the OD task, we investigated whether changes in BOLD activity within the attention network were correlated with behavioral improvements. Specifically, we examined the relationship between beta modulation measured in the DVAN (calculated as the average modulation across all ROIs embedded in the network, per subject) and changes in OD performance, reported in our previous work [7]. A robust correlation analysis (skipped Pearson correlation, see *Methods section* for statistical details) revealed a significant positive correlation (r = 0.465, p = 0.045, 95%, CI [0.097, 0.708], t = 2.164), indicating that participants who showed greater improvement in behavioral performance also exhibited greater modulation in beta activity across the DVAN (*Figure 4a*).

**Figure 4.**
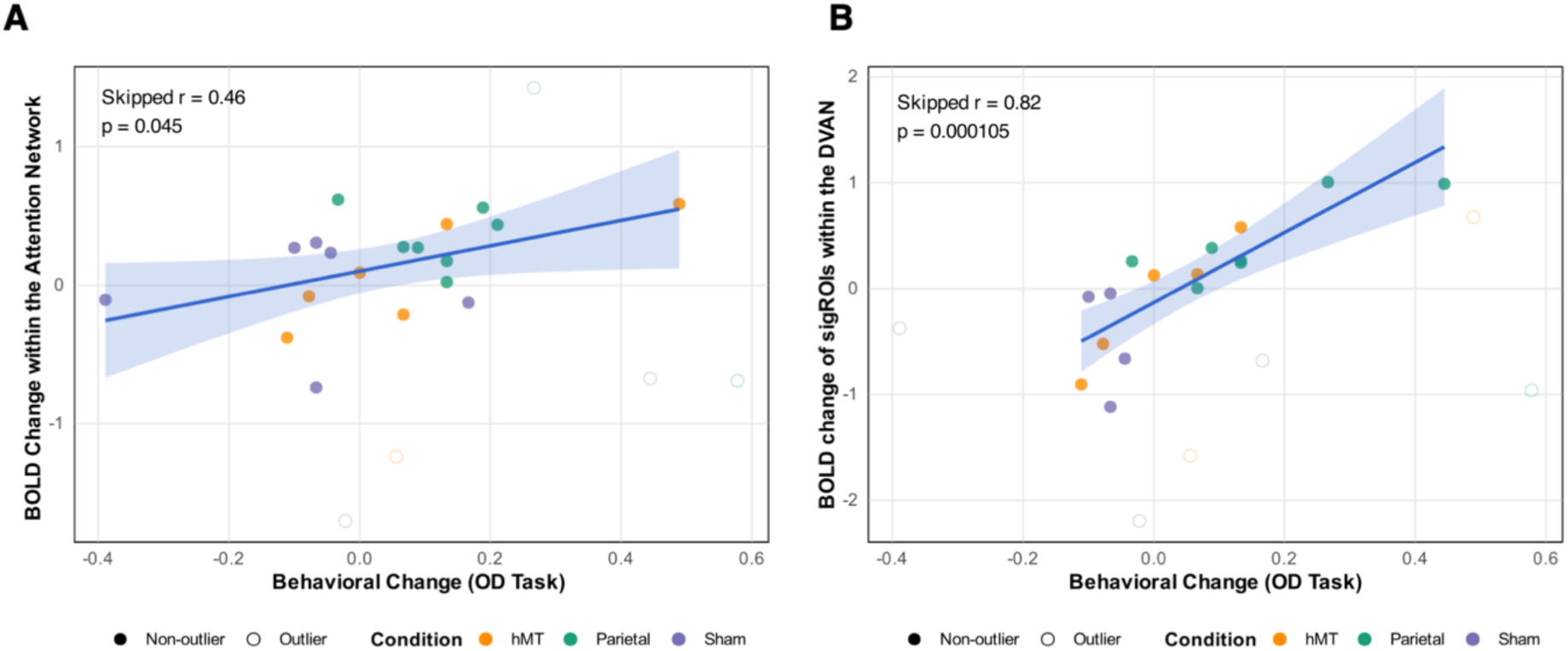
Robust correlation analyses between changes in behavioral performance on the Orientation Discrimination (OD) task and changes in beta values within the Attention Network. (A) Correlation between behavioral changes and beta values across all ROIs embedded in the Attention Network (r = 0.465, p = 0.045). (B) Correlation between behavioral changes and beta values in significant ROIs (r = 0.819, p = 0.0001). Shaded areas represent the 95% bootstrap confidence intervals. Both analyses were conducted using Robust Correlation approach ([45]), which detects and removes bivariate outliers based on robust Mahalanobis distances. Dots represent individual data points (colored per Stimulation Condition); data points identified as outliers were excluded from the analysis.

To further explore the specificity of this effect, we conducted an additional correlation analysis focusing on the ROIs that resulted the most affected by stimulation (hMT, anterior IPS, posterior IPS; as reported in the second part of the Results section). A skipped Pearson correlation (using the same statistical approach as in the previous model) was then employed to correlate the mean change in BOLD signal measured within these brain areas (anterior and poster IPS, FEF, and hMT), with the change in behavioral performance in the OD task. This analysis revealed a stronger positive correlation (r = 0.819, p = 0.0001), with a bootstrap-estimated 95% confidence interval of [0.676, 0.922], and an associated t-statistic of 5.337. These results suggest that the neural changes evoked by stimulation in these brain regions are closely correlated with behavioral improvements (*Figure 4b*).

## Discussion

This study investigated the effects of multi-session transcranial random noise stimulation (tRNS) on learning-related plasticity in the dorsal-ventral attention network (DVAN) during two distinct perceptual tasks. Our results show that parietal tRNS targeting the bilateral intraparietal sulcus (IPS) selectively increased BOLD activity only during an orientation discrimination (OD) task, and that this functional increase correlated with behavioral improvement. This task-selective modulation highlights the interaction between tRNS and task-relevant neural circuits, suggesting that stimulation effects depend on the specific cognitive demands of the task and the functional engagement of targeted regions. Control analyses of the default mode network (DMN) showed no significant changes from pre- to post-training in any of the stimulation conditions, confirming the specificity of stimulation effects to task-relevant networks. These findings indicate that tRNS preferentially amplifies activity in neural circuits that are both highly engaged and optimally tuned to the cognitive demands of a task.

The OD task elicited increased BOLD signal modulation within key nodes of the DVAN, particularly in the anterior and posterior IPS, regions critical for visuospatial attention ([46], [47], [48], [49]). Through a multi-session stimulation protocol targeting these key nodes, tRNS strengthens activity in areas already highly engaged in the task, modulates cortical activity across the network, and facilitates perceptual learning by amplifying neural processing in the most functionally relevant regions. The absence of similar effects during TOJ further supports the hypothesis that tRNS preferentially enhances neural activity in circuits with heightened baseline excitability due to task engagement ([7][50]). Crucially, stimulation of hMT, did not exert the same effects at either the cortical or behavioral level, further emphasizing the importance of targeting critical regions for significant network-wide modulation.

Notably, no significant changes in BOLD signal within the DVAN were observed during the TOJ task across different stimulation conditions. This suggests that tRNS-induced neural modulation is contingent on the task’s cognitive demands and the specific neural substrates involved. One possible explanation is that tRNS amplifies neural activity in regions with heightened baseline excitability due to task engagement, consistent with findings showing that transcranial electrical stimulation (tES) can increase the neural signal-to-noise ratio ([35], [37], [51]). The absence of modulation for the TOJ task may indicate that the neural mechanisms supporting temporal resolution were either less responsive to tRNS or that the task paradigm did not sufficiently engage these networks to elicit detectable changes. This interpretation aligns with evidence suggesting that tES effects are state-dependent, modulated by the baseline activity of the targeted regions and the cognitive context in which stimulation occurs ([52], [53]).

A key finding was the contrasting patterns of neural activity observed after active stimulation versus sham conditions. While parietal tRNS increased BOLD responses within the attention network, the sham condition showed a significant decline in activity without corresponding behavioral improvements (full behavioral results are reported in [7]). This suggests that the observed downregulation is not simply an efficiency-driven adaptation ([22], [28], [54] but may instead reflect premature stabilization of neural activity, limiting further learning. Thus, tRNS appeared to counteract this usually adaptive downregulation by sustaining or increasing neural activity in the attention network regions involved in spatial attention, thus promoting learning ([15]). This suggests that tRNS may prevent the natural tendency of the brain to save on neural resources after learning, leading to sustained plasticity and enhanced cognitive functions. The random fluctuating electrical input of tRNS could play a crucial role in maintaining cortical excitability. Unlike transcranial direct current stimulation (tDCS), which delivers a steady current and may induce neural habituation ([55], [56], [57]), the random nature of tRNS prevents early stabilization, allowing continued engagement of the neural circuits involved in learning and maintaining cortical excitability. These findings provide further evidence that non-invasive neuromodulation techniques can shape plasticity dynamics by modulating cortical excitability patterns over time.

When analyzing the effects of tRNS across different regions of the DVAN, we found that stimulation selectively enhanced activity in specific nodes, including the left hMT, bilateral anterior IPS, and bilateral posterior IPS. This regional specificity suggests that tRNS effects are not uniform across the network but instead concentrate in the most functionally engaged areas ([36], [38], [58], [59]). These findings underscore the parietal response associated with spatial attention ([46], [60], [61], [62]), and further indicate its significant responsiveness to neuromodulation. Also, the absence of significant changes in other DVAN regions reinforces the notion that neuromodulation interventions should be precisely localized to maximize cognitive benefits.

Our study extends previous research on tES-induced BOLD modulation, which has primarily focused on single-session transcranial direct current stimulation (tDCS) or transcranial alternating current stimulation (tACS), with very limited exceptions ([63]). We provide the first evidence of long-term neural modulation induced by multi-session tRNS, demonstrating its potential for sustained enhancement of task-relevant network activity ([40], [64], [65]. Notably, our findings align with resting-state data from our previous study ([7]), which showed increased functional connectivity within the DVAN following parietal tRNS, further supporting the hypothesis that tRNS enhances neural plasticity through targeted and prolonged network engagement.

Finally, the correlation models suggest that changes in behavioral performance on the OD task are significantly related to neural modulation within the Attention Network as a whole. Notably, this relationship is much stronger when considering the most affected ROIs by stimulation, indicating that specific regions within the network may play a more prominent role in driving these behavioral changes.

In summary, by preventing task-induced neural downregulation, tRNS sustains cortical excitability, prolongs plasticity, and enhances cognitive function. These findings highlight its potential for improving visuospatial attention and suggest broader applications for cognitive enhancement in both healthy individuals and clinical populations ([66], [67], [68], [69]). Future research should explore whether optimized tRNS protocols can generalize to more complex cognitive tasks and whether longer or more intense stimulation further amplifies its beneficial effects.

### Conclusions

By demonstrating that tRNS selectively enhances neural activity in the attention network and promotes visuospatial learning, our study highlights the potential of non-invasive stimulation for modulating cortical plasticity. The observed increase in BOLD responses within task-relevant regions suggests that combining tRNS with targeted cognitive training could be a promising strategy for enhancing perceptual learning. These findings have broad implications for developing non-invasive interventions aimed at improving cognitive function in both healthy individuals and clinical populations. Further research is needed to explore optimal stimulation parameters and investigate the applicability of these findings to disorders characterized by attentional deficits.

## Material and Methods

### Participants

Thirty-seven neurologically healthy subjects (20 females, mean age 22.8 years old), with normal or corrected-to-normal vision participated in the study. Rigorous screening ensured that all subjects were free of medical contra-indications for both MRI and brain stimulation. Monetary compensation was provided to all participants in acknowledgement of their commitment to the experimental protocol. Ethical approval was obtained from the University of Trento’s ethical committee. All Subjects were informed about the nature of the study, potential risks, and benefits, and provided written consent before participating.

### Experimental Procedure

Subjects engaged in a comprehensive multi-session experiment that lasted seven days (one session per day; *Figure 4*). On the first session, subjects underwent a staircase procedure that assessed their individual psychophysical thresholds in two attention tasks: an orientation discrimination task (OD), and a temporal order judgment task (TOJ). Both tasks utilized the same visual stimuli consisting of a pair of sine-wave gratings (Gabors, see *Stimuli* section for details). Thresholds for the TOJ and OD tasks were determined separately using 3-1 staircase procedures. Incorrect responses decreased task difficulty, while three consecutive correct responses increased difficulty. The staircases terminated after 30 reversals, and thresholds were calibrated based on these values for the subsequent sessions.

On the second session (pre-test session), participants performed these two attention tasks while in the MRI, with trials of both tasks randomly interleaved within each block. Participants viewed stimuli with fixed temporal offsets (TOJ) and orientation values (OD) set at individual threshold levels determined on the first session. Subjects were cued randomly to attend to either temporal order or orientation feature: each trial began with a 2-second instruction interval where a cue-word (“*Time*“ or “*Orientation*”) indicated the task for the upcoming trial. Following the cue, stimuli were presented for 500 msec, and subjects made forced-choice judgments during a response interval of 1.5 seconds. The task cue was presented again during the response interval, and the trial concluded with a fixation interval of 2 or 4 seconds. Subjects completed a total of five blocks (corresponding to five MRI scans; see *Neuroimaging Procedures* sections for details).

Subsequent sessions three through six constituted the training and stimulation phase. During these sessions, subjects received 25 minutes hf-tRNS or sham stimulation while concurrently training on the same OD and TOJ tasks. The two trial types were randomly interleaved as the task were presented as in the pre-test. On the last session (post-test session), participants replicated the procedure from the pre-test sessions, performing the OD and TOJ tasks while in the MRI. This comprehensive experimental design allowed for the systematic exploration of the effects of hf-tRNS on attentional tasks.

### Stimuli

Visual stimuli employed in the study were identical to those used in our previously published work ([7]), consisting of a pair of sine-wave gratings (Gabors) with fixed temporal offsets and orientation values tailored to individual thresholds. Stimulus presentation and task execution were controlled using Matlab r2016a and Psychtoolbox 3.0.8. For a detailed description of the stimuli used, please refer to the previous work ([7]).

**Figure 4.**
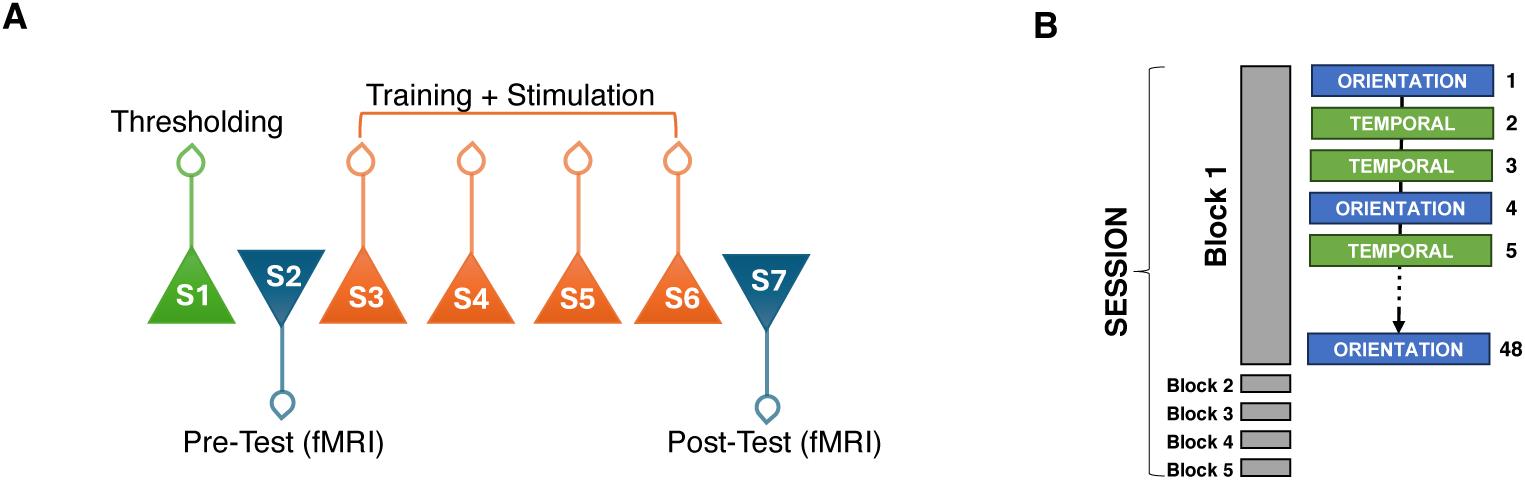
Experimental procedure. **A) Experiment Timeline.** On Day 1 subjects underwent the threshold procedure. On Day 2 subjects were tested on the tasks while fMRI data were collected (pre-test session). From Day 3 to Day 6, subjects underwent behavioral training concurrently with tRNS for 25 minutes. Day 7 was a repeat of Day 2 (post-test session). **B) Session, with blocks and trials sequence example**. One session comprised 5 blocks: in each block two types of attention tasks (OD and TOJ trials, 48 trials per block for a total of 240 per session) were randomly presented within each block.

**Figure5.**
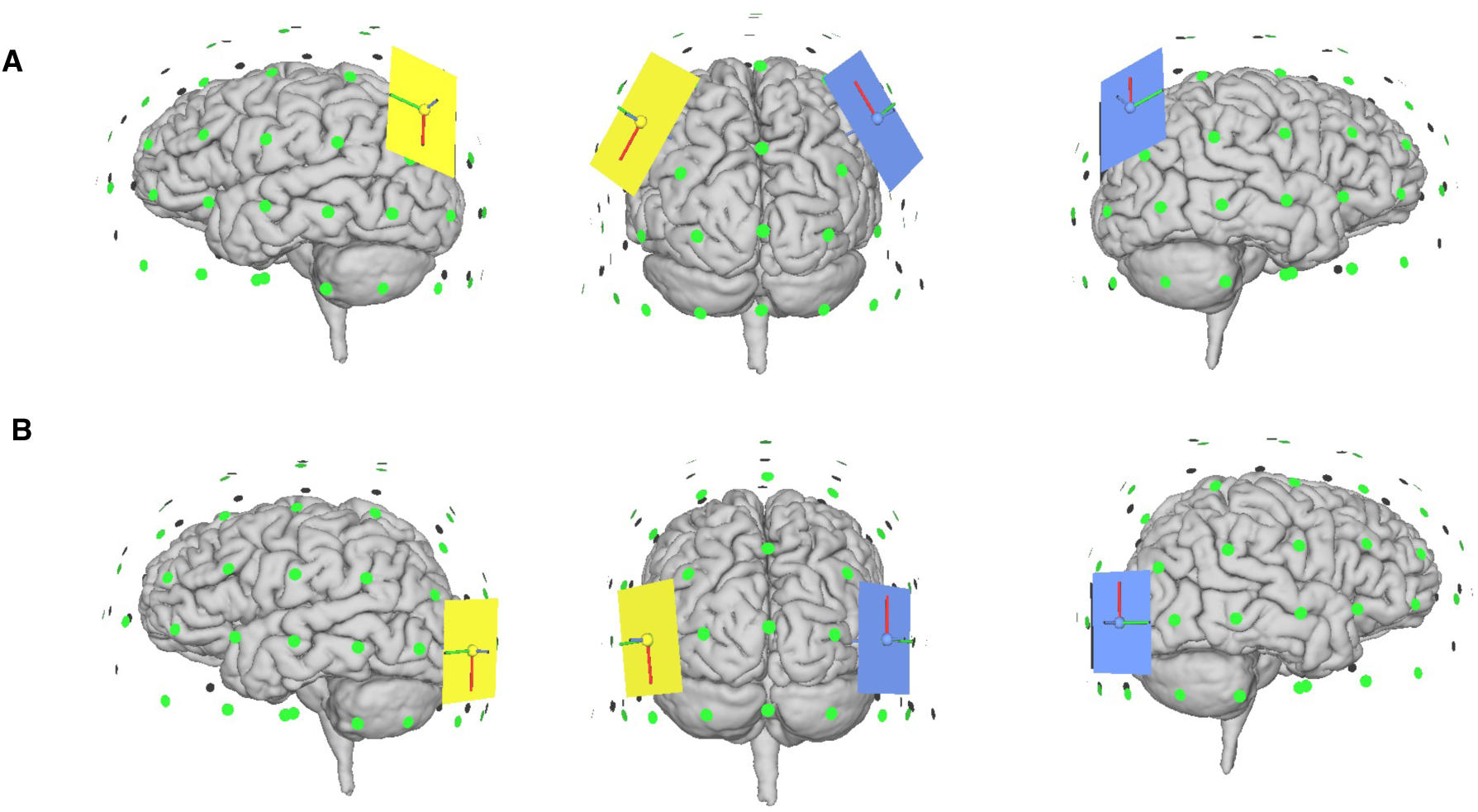
Stimulation Settings. Stimulation sites were localized using the EEG 10/20 system. Electrodes were placed bilaterally over P3 and P4 for Parietal stimulation (Panel A) and for Sham stimulation (Panel A, no active stimulation), and over P07 and P08 for hMT stimulation (Panel B). Images samples were produced using SimNIBS [71], [72]).

### Stimulation Protocol

Participants underwent high-frequency tRNS (hf-tRNS; frequency range 101-640Hz) or sham stimulation targeting specific brain regions, following the international Electroencephalographic 10/20 system for scalp electrode localization ([70]). For hf-tRNS targeting hMT region, electrodes were centered bilaterally over PO7/PO8 (left and right, respectively), while for parietal (IPS) and sham stimulation, electrodes were centered over P3/P4 (*Figure 2*). Stimulation procedure details are described in the previously published work [7].

### Neuroimaging Procedure (Fast event related fMRI protocol)

Brain imaging sessions were conducted at the Center for Mind and Brain Science of the University of Trento, using a 4T Bruker MedSpec MRI scanner equipped with an 8-channel head-coil. Each session began with the acquisition of high-resolution T1-weighted images (MP-RAGE 1×1×1 mm voxel size, 176 sagittal slices) for anatomical reference. Functional imaging comprised T2*-weighted EPIs (TR=2.0 seconds, TE=28.0, flip angle=73∘, 3×3×3mm voxel size, 0.99mm gap, 30 axial slices acquired interleaved, and a 192 mm FOV) were collected during each session. Each task-related functional run consisted of 185 volumes were acquired, while resting-state functional runs consisted of 120 volumes. Each scanning session included one anatomical run followed by six functional runs (five task runs and one resting-state run performed after every three task runs to allow subjects to rest), lasting approximately one hour.

During functional task runs, we used a Fast Event-Related (FER) fMRI protocol, a neuroimaging experimental design that allows for the investigation of brain responses to individual events or stimuli rapidly presented with a short interstimulus interval (ISI) (Dale 1999; Handwerker et al., 2001). Using a FER fMRI design, we investigated changes in BOLD activity within the primary nodes of the DVAN following multi-session tRNS combined with cognitive training. This analysis would further allow us to assess whether the task-related activity exhibits a similar pattern of results as the resting-state data ([7]).

In each run, subjects viewed either a TOJ or OD trial, presented as a sequential pair of Gabors gratings (see Stimuli section for details) via a periscope mirror mounted on the MR head-coil. Each trial lasted 3.75sec and was followed by a variable ISI (ranging from 2 to 4 seconds). These display presentation parameters are the one used in the previous study published ([7]). No feedback was provided. Stimuli were balanced within each run, such that each type of trial appeared an equal number of times in all five runs. Total number of trials per scanning session was 240 (120 per trial type), for a total of 480 trials per subject (240 for the pre-test session, and 240 for the post-test session). After three blocks, a resting-state scan was acquired. The change in the BOLD signal was assessed by measuring the event-related BOLD response changes pre- and post-test sessions.

### Task-related fMRI data analysis

The functional and anatomical data analyses were conducted using BrainVoyager QX (Brain Innovations Inc., Maastricht, The Netherlands). The preprocessing pipeline of the task-related data included the removal of the first 6 volumes to account for non-steady state magnetization (leaving 179 volumes for subsequent analysis), 3D motion correction, slice-timing correction, realignment, and spatial smoothing using a 4mm FWHM Gaussian kernel (full-width at half maximum). Runs with head motion exceeding 3mm in any given dimension were discarded and subjects were eliminated from further analysis when a number equal or greater of 2 runs was excluded in one of the two fMRI sessions (two subjects were eliminated in this step). After preprocessing, the functional scans were initially automatically co-registered and then manually adjusted to each individual’s high-resolution anatomical scan. Following co-registration, images were normalized into standardized Talairach coordinates ([73], [74]). A 3D aligned time courses (VTCs) was created for each run after the intra-session anatomical was aligned with the high resolution anatomical.

Next, single-subject deconvolution analysis was performed. We used a deconvolution analysis to analyze the BOLD responses between different stimuli conditions. Beta-weight values over 20-second period following stimulus onset were contrasted within the target processing region. The individual mapping data were subjected to a standard general linear model (GLM) with one predictor for each condition (TOJ and OD) against a rest condition. We then analyzed the deconvolved BOLD response in 19 ROIs included in the DVAN: the right/left anterior, posterior and medial IPS, right/left hMT, right and left frontal eye fields (FEFs) of the dorsal network; the right/left of the posterior, the right anterior and left/right central temporal parietal junction (TPJ), the right/left ventral frontal cortex (VFC), and the right/left inferior frontal junction (IFJ) of the ventral networks. These regions were identified using standardized mean coordinates (*Table 2*) derived from the literature ([8], [39], [75], [76]). Relative to our previous work that investigated functional connectivity within the DVAN ([7]), we included additional ROIs (e.g., IPS was subdivided into posterior, anterior and medial IPS) to better cover the change in activation of the attention network. In fact, this subdivision was important because each region plays a distinct role in attentional processes: the posterior IPS is primarily involved in spatial attention and the integration of visual information ([77]) , while the anterior IPS is associated with the allocation of attentional resources and task relevance ([78]) and the medial IPS is involved in visuospatial attention ([79], [80]). By capturing these differences, we can better understand the varying activation patterns within the attention network and their relationship to behavioral outcomes.

To control for network-specific effects of stimulation, we also investigated whether tRNS and training influenced cortical activity within a different control network, the default mode network (DMN), which is typically active during passive or resting conditions ([44], [81]). The same procedure used for the DVAN analysis was employed to extract the time series and analyze cortical activity within 10 main ROIs of the DMN (same ROIs used in the functional connectivity analysis of the previous work; [7]). These included: the ventral medial prefrontal cortex (vMPFC), the dorsal prefrontal cortex (dPFC), the left posterior cingulate cortex (PCC), the left lateral temporal cortex (TPC), the bilateral inferior parietal lobe (IPL), the bilateral parahippocampal cortex (PHC), and bilateral hippocampal formation (HF+). Coordinates in MNI space were mapped onto Talairach space using the ‘MNI-to-Talairach converter with Brodmann Areas tool’ included in the Yale BioImage Suite Package ([82]).

Functional data from all voxels within a 4-mm radius sphere from the center coordinate of each ROI were averaged into a single time series, keeping the ROIs similar in size across participants. The time courses were then extracted from each ROI and an individual GLM was applied to estimate the response amplitude, independently for each subject. The resulting beta weights estimate peak activation for each run, assuming a standard 2γ model of the hemodynamic response function. These estimates (peak of the beta weights) from each individual ROI were used as the input to the subsequent statistical analysis.

**Figure 6.**
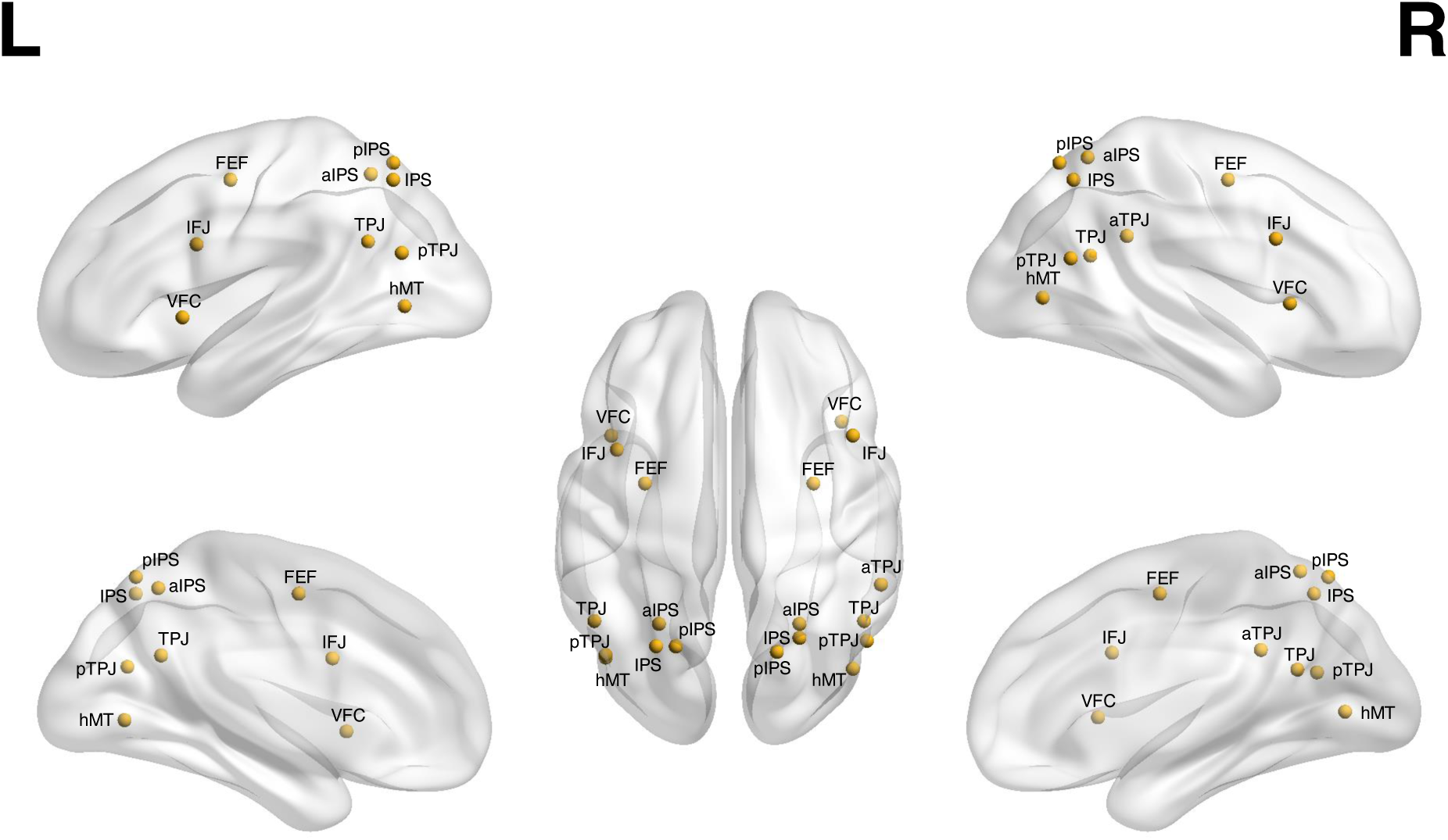
Brain representation of the ROIs embedded in the attention network. Three-dimensioned views (sagittal and coronal views) of the ROI spheres embedded in the attention network, visualized with BrainNet Viewer package for visualization ([83]).

### Data Analysis

We conducted an analysis of BOLD signal changes in response to attentional tasks, specifically the OD and the TOJ task. Peaks of the beta weights relative to individual ROI collected during pre- and post-stimulation fMRI session (pre- and post-sessions) were analyzed. To evaluate the impact of stimulation combined with training on the attentional network, we calculated beta weights representing the overall modulation of the DVAN. These beta weights were derived within each ROI first and then averaged across ROIs to obtain a single beta weight per subject, reflecting the overall trend in BOLD signal modulation ([84]).

First, to assess baseline comparability between stimulation groups prior to stimulation (at baseline), we conducted a linear mixed-effects model using *lmer*() function from the lme4 package ([85]) in RStudio (version 2023.06.0). Next, delta scores representing the difference between pre- and post-stimulation beta weights were computed on individual subject (Δβ = β(fMRI post-test)– β(fMRI pre-test) to quantify changes in BOLD signal modulation. These delta measures reflect the stimulation-induced changes in BOLD signal normalized to baseline. Within-subject delta beta weights were derived to capture the individualized BOLD modulation rather than a group average approach ([86], [87]; for further discussion about differences between within-subjects versus group-averaged approaches, see [88]). A mixed effects model was then fit on subjects’ delta beta weights, where the between-subjects predictor was Stimulation (Parietal, hMT, Sham), and the within-subjects predictor was Task (OD and TOJ). Model comparisons were conducted, and the selected model formula was *lmer*(Δβ ∼ Stimulation*Task+(1+session|subject), data = df_delta_betas). This linear mixed-effects model was based on the delta scores, and random intercepts and slopes were included for each subject. Chi-squared and p-values were reported for both interactions and main effects. After fitting a linear mixed-effects model, estimated marginal means (EMMs) were computed for each level of the stimulation factor. Contrasts between stimulation levels were then estimated, and associated standard errors, confidence intervals, t-ratios, and p-values were calculated. Adjustments for multiple comparisons were made using the FDR method ([89]). The same statistical analyses were then computed on the beta weights of the main ROIs included in DMN to examine whether stimulation and training induced changes in cortical activity in this control network.

Next, we compared the change in beta values measured within the DVAN and the DMN. A mixed-effects model was fitted on subjects’ mean delta beta scores of both networks, with network (DVAN and DMN) and task (OD and TOJ) serving as the within-subject predictor and stimulation (Parietal, hMT, Sham) as the between-subjects predictor. This model served to evaluate whether there was a difference in signal modulation within the two networks depending on stimulation conditions, hence indicating whether stimulation affected a specific network or was general in the whole brain.

Finally, we examine the relationship between changes in behavioral performance and modulation of BOLD activity, we employed a skipped Pearson correlation ([90], [91], [92]). This robust statistical approach is designed to detect and remove bivariate outliers based on Mahalanobis distances, ensuring that the correlation estimate is not disproportionately influenced by extreme values. This method was applied to investigate the association between changes in beta values across ROIs in the Attention Network (averaged per subject) and improvements in OD task performance. The significance of the correlation was assessed using bootstrap resampling, which provides confidence intervals that account for data variability and potential deviations from normality. To further explore the specificity of the observed effect, we conducted an additional correlation analysis, focusing on the ROIs most affected by stimulation (*hMT, anterior IPS, and posterior IPS*). The beta modulation of these ROIs was averaged for each participant and correlated with OD performance changes. Only participants who were included in both the task-related BOLD analysis and the behavioral analysis (i.e., those not excluded due to motion or other factors) were considered in this analysis. All statistical analyses were conducted using MATLAB (statistics) and Rstudio (Figures), with robust statistical functions from the Robust Correlation Toolbox ([93], [94]).

## Appendix

**Table 1 Appendix.** Results of the contrasts between Stimulation conditions for each ROI divided per hemisphere embedded in the DVAN relative to the OD task. For each level of brain area (ROI), statistical details are reported comprising estimated marginal means for each combination of Stimulation factor, along with SE, t ratio and corresponding p value (FDR-corrected).

| ROI | Left |  |  |  |  |  | Right |  |  |  |  |  |
| --- | --- | --- | --- | --- | --- | --- | --- | --- | --- | --- | --- | --- |
|  | Contrast | Estimate | SE | df | t.ratio | p.value | Contrast | Estimate | SE | Df | t.ratio | p.value |
| FEF | hMT - Parietal | -0.2713 | 0.416 | 401 | -0.652 | 0.8696 | hMT - Parietal | -0.4496 | 0.416 | 401 | -1.081 | 0.4208 |
|  | hMT - Sham | -0.0358 | 0.416 | 401 | -0.086 | 0.9316 | hMT - Sham | 0.1608 | 0.416 | 401 | 0.386 | 0.6994 |
|  | Parietal vs Sham | 0.2356 | 0.425 | 401 | 0.554 | 0.8696 | Parietal vs Sham | 0.6104 | 0.425 | 401 | 1.436 | 0.4208 |
| hMT | hMT - Parietal | -0.6628 | 0.416 | 401 | -1.593 | 0.1119 | hMT - Parietal | -0.1174 | 0.416 | 401 | -0.282 | 0.7780 |
|  | hMT - Sham | 0.7152 | 0.416 | 401 | 1.719 | 0.1119 | hMT - Sham | 0.5865 | 0.416 | 401 | 1.410 | 0.2392 |
|  | Parietal vs Sham | 1.3780 | 0.425 | 401 | 3.242 | 0.0039 | Parietal vs Sham | 0.7038 | 0.425 | 401 | 1.656 | 0.2392 |
| IFJ | hMT - Parietal | -0.4338 | 0.416 | 401 | -1.043 | 0.7344 | hMT - Parietal | -0.4135 | 0.416 | 401 | -0.994 | 0.4814 |
|  | hMT - Sham | -0.1399 | 0.416 | 401 | -0.336 | 0.7369 | hMT - Sham | 0.0366 | 0.416 | 401 | 0.088 | 0.9300 |
|  | Parietal vs Sham | 0.2939 | 0.425 | 401 | 0.692 | 0.7344 | Parietal vs Sham | 0.4501 | 0.425 | 401 | 1.059 | 0.4814 |
| IPS | hMT - Parietal | -0.7819 | 0.416 | 401 | -1.879 | 0.1828 | hMT - Parietal | 0.0353 | 0.416 | 401 | 0.085 | 0.9323 |
|  | hMT - Sham | -0.2211 | 0.416 | 401 | -0.531 | 0.5954 | hMT - Sham | 0.8859 | 0.416 | 401 | 2.129 | 0.0691 |
|  | Parietal vs Sham | 0.5608 | 0.425 | 401 | 1.320 | 0.2816 | Parietal vs Sham | 0.8505 | 0.425 | 401 | 2.001 | 0.0691 |
| Ant IPS | hMT - Parietal | -1.0075 | 0.416 | 401 | -2.422 | 0.0238 | hMT - Parietal | -0.3062 | 0.416 | 401 | -0.736 | 0.4621 |
|  | hMT - Sham | 0.0541 | 0.416 | 401 | 0.130 | 0.8965 | hMT - Sham | 0.8762 | 0.416 | 401 | 2.106 | 0.0537 |
|  | Parietal - Sham | 1.0617 | 0.425 | 401 | 2.498 | 0.0238 | Parietal vs Sham | 1.1824 | 0.425 | 401 | 2.782 | 0.0170 |
| Post IPS | hMT - Parietal | -0.4238 | 0.416 | 401 | -1.019 | 0.3090 | hMT - Parietal | -0.7931 | 0.416 | 401 | -1.906 | 0.0860 |
|  | hMT - Sham | 1.0413 | 0.416 | 401 | 2.503 | 0.0191 | hMT - Sham | 0.4753 | 0.416 | 401 | 1.142 | 0.2540 |
|  | Parietal vs Sham | 1.4651 | 0.425 | 401 | 3.447 | 0.0019 | Parietal vs Sham | 1.2684 | 0.425 | 401 | 2.984 | 0.0090 |
| TPJ | hMT - Parietal | -0.4360 | 0.416 | 401 | -1.048 | 0.5564 | hMT - Parietal | -0.4704 | 0.416 | 401 | -1.131 | 0.4484 |
|  | hMT - Sham | -0.0553 | 0.416 | 401 | -0.133 | 0.8943 | hMT - Sham | -0.0284 | 0.416 | 401 | -0.068 | 0.9457 |
|  | Parietal vs Sham | 0.3807 | 0.425 | 401 | 0.896 | 0.5564 | Parietal vs Sham | 0.4420 | 0.425 | 401 | 1.040 | 0.4484 |
| Ant TPJ | hMT - Parietal | - | - | - | - | - | hMT - Parietal | 0.1454 | 0.416 | 401 | 0.350 | 0.7906 |
|  | hMT - Sham | - | - | - | - | - | hMT - Sham | 0.2584 | 0.416 | 401 | 0.621 | 0.7906 |
|  | Parietal vs Sham | - | - | - | - | - | Parietal vs Sham | 0.1129 | 0.425 | 401 | 0.266 | 0.7906 |
| Post TPJ | hMT - Parietal | -0.0971 | 0.416 | 401 | -0.233 | 0.9508 | hMT - Parietal | -0.6807 | 0.416 | 401 | -1.636 | 0.1539 |
|  | hMT - Sham | -0.1233 | 0.416 | 401 | -0.296 | 0.9508 | hMT - Sham | 0.0427 | 0.416 | 401 | 0.103 | 0.9184 |
|  | Parietal vs Sham | -0.0262 | 0.425 | 401 | -0.062 | 0.9508 | Parietal vs Sham | 0.7234 | 0.425 | 401 | 1.702 | 0.1539 |
| VFC | hMT - Parietal | -0.5409 | 0.416 | 401 | -1.300 | 0.2914 | hMT - Parietal | -0.6973 | 0.416 | 401 | -1.676 | 0.2354 |
|  | hMT - Sham | 0.0478 | 0.416 | 401 | 0.115 | 0.9085 | hMT - Sham | -0.0946 | 0.416 | 401 | -0.227 | 0.8202 |
|  | Parietal vs Sham | 0.5888 | 0.425 | 401 | 1.385 | 0.2914 | Parietal vs Sham | 0.6027 | 0.425 | 401 | 1.418 | 0.2354 |

**Table 2 Appendix.** Results of the LMM for each ROI embedded in the DVAN, with data collapsed across hemispheres for the TOJ task. For each brain area (left and right hemisphere data were collapsed), statistical details of the LMM fitted (lmer(normalized_delta ∼ Condition + (1|subj), data=df_ROI) are reported comprising of Chi-square and corresponding p-value of the model.

| ROI | ChiSq | Df | Pr(>Chisq) |
| --- | --- | --- | --- |
| FEF | 3.0279 | 2 | 0.2200 |
| hMT | 8.2966 | 2 | 0.0158 |
| IFJ | 3.7083 | 2 | 0.1566 |
| IPS | 1.4832 | 2 | 0.4764 |
| Ant IPS | 1.0495 | 2 | 0.5917 |
| Post IPS | 1.3139 | 2 | 0.5184 |
| TPJ | 3.8990 | 2 | 0.1423 |
| Post TPJ | 0.5534 | 2 | 0.7583 |
| VFC | 2.5490 | 2 | 0.2796 |

**Table 3 Appendix.** Results of the contrasts between Stimulation conditions for each ROI divided per hemisphere embedded in the DVAN relative to the TOJ task. For each level of brain area (ROI), statistical details are reported comprising estimated marginal means for each combination of Stimulation factor, along

| ROI | Left |  |  |  |  |  | Right |  |  |  |  |  |
| --- | --- | --- | --- | --- | --- | --- | --- | --- | --- | --- | --- | --- |
|  | Contrast | Estimate | SE | df | t.ratio | p.value | Contrast | Estimate | SE | Df | t.ratio | p.value |
| FEF | hMT - Parietal | -0.2639 | 0.427 | 386 | -0.618 | 0.8359 | hMT - Parietal | -0.2735 | 0.427 | 386 | -0.641 | 0.9070 |
|  | hMT - Sham | -0.0885 | 0.427 | 386 | -0.207 | 0.8359 | hMT - Sham | -0.0499 | 0.427 | 386 | -0.117 | 0.9070 |
|  | Parietal vs Sham | 0.1754 | 0.436 | 386 | 0.402 | 0.8359 | Parietal vs Sham | 0.2235 | 0.436 | 386 | 0.513 | 0.9070 |
| hMT | hMT - Parietal | -0.3496 | 0.427 | 386 | -0.819 | 0.6200 | hMT - Parietal | -0.5596 | 0.427 | 386 | -1.311 | 0.5721 |
|  | hMT - Sham | 0.1611 | 0.427 | 386 | 0.377 | 0.7061 | hMT - Sham | -0.3565 | 0.427 | 386 | -0.835 | 0.6063 |
|  | Parietal vs Sham | 0.5107 | 0.436 | 386 | 1.171 | 0.6200 | Parietal vs Sham | 0.2031 | 0.436 | 386 | 0.466 | 0.6416 |
| IFJ | hMT - Parietal | -0.5596 | 0.427 | 386 | -1.311 | 0.5721 | hMT - Parietal | -0.4761 | 0.427 | 386 | -1.115 | 0.6306 |
|  | hMT - Sham | -0.3565 | 0.427 | 386 | -0.835 | 0.6063 | hMT - Sham | -0.2662 | 0.427 | 386 | -0.623 | 0.6306 |
|  | Parietal vs Sham | 0.2031 | 0.436 | 386 | 0.466 | 0.6416 | Parietal vs Sham | 0.2099 | 0.436 | 386 | 0.481 | 0.6306 |
| IPS | hMT - Parietal | -0.6729 | 0.427 | 386 | -1.576 | 0.1737 | hMT - Parietal | -0.0767 | 0.427 | 386 | -0.180 | 0.8575 |
|  | hMT - Sham | -0.9645 | 0.427 | 386 | -2.259 | 0.0733 | hMT - Sham | 0.3448 | 0.427 | 386 | 0.808 | 0.6298 |
|  | Parietal vs Sham | -0.2916 | 0.436 | 386 | -0.669 | 0.5041 | Parietal vs Sham | 0.4215 | 0.436 | 386 | 0.966 | 0.6298 |
| Ant IPS | hMT - Parietal | -0.8303 | 0.427 | 386 | -1.945 | 0.1575 | hMT - Parietal | 0.3653 | 0.427 | 386 | 0.856 | 0.5891 |
|  | hMT - Sham | -0.1777 | 0.427 | 386 | -0.416 | 0.6775 | hMT - Sham | 0.3915 | 0.427 | 386 | 0.917 | 0.5891 |
|  | Parietal vs Sham | 0.6526 | 0.436 | 386 | 1.497 | 0.2030 | Parietal vs Sham | 0.0263 | 0.436 | 386 | 0.060 | 0.9520 |
| Post IPS | hMT - Parietal | -0.2552 | 0.427 | 386 | -0.598 | 0.5503 | hMT - Parietal | -0.0426 | 0.427 | 386 | -0.100 | 0.9205 |
|  | hMT - Sham | 0.3007 | 0.427 | 386 | 0.704 | 0.5503 | hMT - Sham | 0.2478 | 0.427 | 386 | 0.581 | 0.8428 |
|  | Parietal vs Sham | 0.5560 | 0.436 | 386 | 1.275 | 0.5503 | Parietal vs Sham | 0.2905 | 0.436 | 386 | 0.666 | 0.8428 |
| TPJ | hMT - Parietal | -0.1764 | 0.427 | 386 | -0.413 | 0.6796 | hMT - Parietal | -0.0697 | 0.427 | 386 | -0.163 | 0.8704 |
|  | hMT - Sham | -0.3965 | 0.427 | 386 | -0.929 | 0.6796 | hMT - Sham | -0.6026 | 0.427 | 386 | -1.411 | 0.3337 |
|  | Parietal vs Sham | -0.2200 | 0.436 | 386 | -0.505 | 0.6796 | Parietal vs Sham | -0.5329 | 0.436 | 386 | -1.222 | 0.3337 |
| Ant TPJ | hMT - Parietal | - | - | - | - | - | hMT - Parietal | 0.3594 | 0.427 | 386 | 0.842 | 0.6005 |
|  | hMT - Sham | - | - | - | - | - | hMT - Sham | -0.0351 | 0.427 | 386 | -0.082 | 0.9345 |
|  | Parietal vs Sham | - | - | - | - | - | Parietal vs Sham | -0.3946 | 0.436 | 386 | -0.905 | 0.6005 |
| Post TPJ | hMT - Parietal | 0.0472 | 0.427 | 386 | 0.111 | 0.9120 | hMT - Parietal | -0.1401 | 0.427 | 386 | -0.328 | 0.7430 |
|  | hMT - Sham | -0.1639 | 0.427 | 386 | -0.384 | 0.9120 | hMT - Sham | -0.3066 | 0.427 | 386 | -0.718 | 0.7430 |
|  | Parietal vs Sham | -0.2111 | 0.436 | 386 | -0.484 | 0.9120 | Parietal vs Sham | -0.1665 | 0.436 | 386 | -0.382 | 0.7430 |
| VFC | hMT - Parietal | -0.4765 | 0.427 | 386 | -1.116 | 0.5917 | hMT - Parietal | -0.5900 | 0.427 | 386 | -1.382 | 0.5034 |
|  | hMT - Sham | -0.2424 | 0.427 | 386 | -0.568 | 0.5917 | hMT - Sham | -0.4092 | 0.427 | 386 | -0.959 | 0.5075 |
|  | Parietal vs Sham | 0.2341 | 0.436 | 386 | 0.537 | 0.5917 | Parietal vs Sham | 0.1807 | 0.436 | 386 | 0.414 | 0.6788 |

## Notes

### Competing Interest Statement

The authors have declared no competing interest.

### Summary of Updates

A correlation Analysis has been added

